# ImmunoCAR Atlas: a pan-immune single-cell resource for dissecting CAR-T therapy response across patient cohorts

**DOI:** 10.64898/2026.11.23.768526

**Authors:** Kirti Jain, Tanishqua Neeraj Sharma, Anas Ansari, Diksha Singh Rathore, Simran Jit, Radul R. Dev, Sherry Bhalla

**Author notes:** **Correspondence:** Dr. Sherry Bhalla, CSIR-IGIB, New Delhi.

## Abstract

Chimeric antigen receptor (CAR) T-cell therapy has transformed treatment for relapsed/refractory hematologic malignancies, yet 40–60% of patients relapse within one to two years, and the immune correlates separating durable responders from non-responders remain poorly defined. Single-cell transcriptomics has been widely applied to CAR-T cohorts, but the resulting data remain fragmented across small, underpowered studies with inconsistent metadata, tissue sources, sampling timepoints, and annotation schemes, and existing immune atlases each address only part of this problem. Here we present ImmunoCAR Atlas (https://car-t-atlas.igib.res.in/), a harmonized, patient-stratified single-cell resource comprising 5.7 million cells from 1,085 samples and 584 clinically annotated patients, spanning three cancer indications and four FDA-approved CAR-T products, integrated into 16 scVI-corrected reference atlases with patient-level responder/non-responder annotation. Unlike prior resources, ImmunoCAR Atlas preserves the full pan-immune tumor microenvironment, links longitudinal pre- and post-infusion samples to standardized clinical outcomes, and provides an interactive interface with dedicated analytical modules for exploring patient and cellular composition, cell-population shifts, differential gene expression, and cell-cell communication between responders and non-responders. The atlas features four purpose-built tools: metaCIDER (Meta-analysis of Cross-study Immune Differentially Expressed Response genes), which identifies genes consistently associated with response or resistance across datasets; CellMARK (Cross-Atlas Cell Marker Discovery), which nominates reproducible cell-type markers distinguishing response groups; CAR-T RSS (CAR-T Response Signature Score), which validates metaCIDER-nominated or user-supplied gene signatures; and a Gene Search module, which allows users to query genes of interest and explore their associations with responder and non-responder groups across datasets. To our knowledge, ImmunoCAR Atlas is the first harmonized single-cell CAR-T resource to integrate clinical annotations, response-centric analytical modules, and downloadable reference atlases in a single platform facilitating mechanistic studies and the discovery of robust, reproducible biomarkers of CAR-T response and resistance across the treatment continuum.

## Introduction

Chimeric antigen receptor (CAR) T-cell therapy has transformed the treatment of relapsed or refractory haematological malignancies, with CD19- and B-cell maturation antigen (BCMA)-directed products now established across B-cell acute lymphoblastic leukaemia (B-ALL), large B-cell lymphoma (LBCL) and multiple myeloma [1–5]. However, a substantial proportion of patients fail to respond. In the pivotal ZUMA-1 and JULIET trials, objective response rates were 82% with axicabtagene ciloleucel (axi-cel) and 52% with tisagenlecleucel (tisa-cel), respectively [2, 3]. This variability is also evident in real-world populations: analysis of the French DESCAR-T registry reported best overall response rates of 80% for axi-cel and 66% for tisa-cel [6]. Together, these studies show that a sizable fraction of patients do not respond to CAR-T therapy, emphasizing the need to understand the cellular features associated with response and non-response.

Single-cell profiling has provided an increasingly detailed view of the cellular features associated with CAR-T response across different haematological malignancies, CAR-T products and treatment time points. In LBCL, scRNA-seq studies of axi-cel and tisa-cel have profiled infusion products together with longitudinal peripheral blood samples and identified response-associated differences in CAR-T differentiation, memory and regulatory states, as well as in the endogenous immune compartment [7–9]. In B-ALL, single-cell analysis of infusion products has linked early-memory and antigen-specific T-cell states to clinical outcome [10]. BCMA-directed CAR-T studies in multiple myeloma have extended such analyses to longitudinal peripheral blood and bone marrow, revealing treatment-associated changes across CAR-T cells, endogenous T cells and other components of the bone-marrow immune microenvironment [11–13]. Paired scRNA-seq and scTCR-seq further allow transcriptional cell states to be related to clonal expansion and dynamics [8, 14]. However, these datasets have largely been analysed as individual cohorts using study-specific cell annotations, clinical metadata structures, sampling schemes and response classifications. Consequently, comparing the same cellular state or response-associated signature across independent studies, diseases, CAR-T products and treatment time points requires substantial reprocessing and harmonization. A unified resource that standardizes cell identities and clinical metadata while preserving study, tissue, treatment and time-point information would enable these datasets to be interrogated within a common analytical framework.

Several large-scale single-cell resources provide reference maps of immune and tumour ecosystems across tissues and cancer types, including TISCH2 and scImmOmics [15, 16]. TISCH2 integrates single-cell transcriptomic datasets from the tumour microenvironment across multiple cancer types, while scImmOmics provides a manually curated collection of single-cell immune multi-omics datasets. These resources, however, were not designed around CAR-T therapy and therefore do not organize data according to CAR-T product, treatment time point and harmonized clinical response. CAR-T-specific resources have begun to address parts of this need. The Open CAR-T Atlas integrates more than one million cells from 13 studies and enables interrogation of CAR-T-cell phenotypes in relation to clinical outcome [17], while CART-GPT was developed using an atlas of 1.12 million CAR-T-cell transcriptomes from 255 individuals to model treatment response and neurotoxicity [18]. These resources predominantly centre on CAR-T/T-cell populations, leaving the endogenous immune compartment less systematically represented. In parallel, repertoire resources such as TCRdb, VDJdb and McPAS-TCR catalogue TCR sequences, clonotypes and antigen associations [19–21], but are not designed to jointly interrogate clonotype, single-cell transcriptional state and patient-level CAR-T outcome. Thus, the remaining computational need is not simply aggregation of additional single-cell datasets, but a framework in which CAR-T studies can be compared using consistent cell identities, clinical variables and treatment-related metadata while retaining their original study context.

To address this need, we developed **ImmunoCAR Atlas**, a harmonized, patient-stratified resource comprising 5.7 million cells from 1,085 samples and 584 clinically annotated patients across three cancer types and four FDA-approved CAR-T products. The data are organized into 16 scVI-integrated reference atlases, with clinical metadata harmonized at the patient and sample levels, including disease, CAR-T product, tissue, treatment time point and responder/non-responder status. Rather than restricting analysis to CAR-T cells, ImmunoCAR Atlas retains the broader immune compartment, enabling response-associated states to be examined across T, NK, B and myeloid populations. Cell identities are standardized using popV, with additional high-resolution annotation of T-cell states, while the original author-provided annotations are retained alongside the standardized labels to preserve study-specific biological context and annotation provenance. Where available, paired scTCR-seq data are linked at single-cell resolution to transcriptional state, sample, patient and clinical outcome. The interactive interface enables users to query and compare cell populations, transcriptional states and clonotypes across studies, diseases, CAR-T products, tissues, treatment time points and clinical response. ImmunoCAR Atlas therefore provides a common computational framework for evaluating whether cellular and clonal features associated with CAR-T response are reproducible across independent cohorts or specific to particular diseases, products or treatment contexts.

## Materials and Methods

### Data Collection and Curation

Publicly available CAR-T scRNA-seq datasets (n = 43) were identified through a PubMed/GEO literature search up to December 2025. Clinical and experimental metadata were manually curated, and response labels were assigned at the patient level according to each study’s criteria. Unlabelled samples were excluded from downstream analyses. Detailed literature search strategies and study inclusion criteria are provided in Supplementary Methods S1 and S2, respectively. Curated metadata are browsable via Atlas’s "Browse by Study" module.

### Data Processing and Cell Annotation

Each dataset underwent dataset-specific quality control, with cell- and gene-level filtering thresholds applied according to the criteria reported in the corresponding original publication and documented in the ImmunoCAR Atlas Help page (Section 22.33.1). Raw count matrices were normalized using scran (v1.22.1) [22], while datasets available only as normalized matrices were used as provided. Preprocessing was implemented in Scanpy (v1.9.3) [23] in Python (3.11.12). Cell types were annotated using PopV (v0.4.2) [24] with Tabula Sapiens v2 references for raw-count datasets, or CellTypist (v1.6.3) [25] for normalized-only datasets. Study-reported labels were retained where available. T-cell functional states were further characterized using complementary marker-based (STCAT) and transcriptional program–based (StarCAT) approaches. Detailed preprocessing, cell annotation, and T-cell state annotation procedures are provided in Supplementary Sections S3 and S4, respectively.

### Clinical Outcome Harmonization

Study-specific response criteria (e.g., IMWG [26], Lugano [27]) were harmonized into binary responder/non-responder categories using a predefined mapping (details in supplementary methods S5).

### Data Integration

Datasets were integrated using scVI (v1.6.6) [28, 29], with patient ID specified as the batch covariate. The resulting latent representations were used for neighborhood graph construction, dimensionality reduction, and graph-based clustering. Integration quality was evaluated using scIB (v1.1.7) [30], considering batch-removal and biological-conservation metrics combined into a weighted overall score. Details of the scVI configuration, clustering strategy, and individual scIB metrics are provided in Supplementary Methods S6.

### sc-TCR Analysis

Paired TRA/TRB clonotypes were defined from CDR3 sequences (details in Supplementary Methods S7), classified by expansion frequency (rare–hyperexpanded), and compared between response groups (Mann–Whitney U) implemented in SciPy (v1.11.4).

### Compositional analysis

Cell-type composition differences between response groups were assessed per tissue/timepoint using scCODA [31], with patient number (and dataset, for the integrated atlas) as covariates. T-cell subset composition was assessed descriptively (boxplots) given low cell counts. Detailed model specifications, statistical criteria, and analysis procedures are provided in Supplementary Methods S8.

### Cell-Cell Interaction Analysis

Ligand–receptor interactions were inferred per sample with CellPhoneDB (v5.0.1) [32] and compared between responders and non-responders using a directional enrichment score and Wilcoxon rank-sum testing as explained in detail in Supplementary Methods S9.

### Patient Cell Composition

Per-patient, per-timepoint cell-type proportions were computed (details in Supplementary Methods S10); pairwise patient differences were tested by χ² test with Pearson residuals identifying contributing cell types.

### Gene Explorer

Precomputed UMAP embeddings allow gene expression and metadata (clinical, technical, annotation) to be overlaid at the dataset and integrated-atlas level. Further details are provided in Supplementary Methods S11.

### Differential Expression and Pathway Analysis

Differential gene expression was performed on single-cell and patient level with limma-trend [33, 34] on scVI-normalized expression (single dataset) [28, 29, 35], adjusting for patient ID (batch), after removing mitochondrial, ribosomal, and sex-linked genes. DEGs were defined as |logFC| > 0.5 with adj. p < 0.05 (single-cell) or nominal p < 0.05 (pseudobulk, given reduced power after patient-level aggregation), and tested for over-representation (Enrichr) separately for up- and down-regulated genes. The detailed gene filtering criteria, statistical models, pathway libraries and enrichment thresholds are provided in the Supplementary Methods S12-13.

### metaCIDER cross-cohort framework

Per-dataset single-cell and pseudobulk DEGs and enriched pathways meeting fixed thresholds were intersected across ≥50% of datasets to define consensus signals; a random-effects meta-analysis (REML) [36] pooled logFC across datasets. Further details are provided in Supplementary Methods S14-15.

### Pre-infusion Response Signature Identification in DLBCL

Seven pre-treatment DLBCL PBMC datasets profiling anti-CD19 CAR-T-treated patients were initially available for this cross-cohort framework. For each dataset, differential expression between NR and R was computed at two levels: (i) at the single-cell level and (ii) at the pseudobulk level as described in the Supplementary Methods S12-13. One dataset, GSE243325, was excluded from meta-analysis discovery because the bulk RNA-seq dataset from the same study was used for external validation; retaining it would have introduced circularity into the analysis. The remaining six cohorts were carried forward.

REML random-effects meta-analysis was performed independently on the single-cell and pseudobulk DEG outputs, pooling log₂FC estimates and reconstructed standard errors across CD4 and CD8 T cell specific genes. For each gene–cell-type combination, only genes present in all six datasets with a directionally consistent effect (all six datasets showing the same sign of log₂FC) were considered. Significance thresholds were set at nominal p < 0.05 for the single-cell meta-analysis and pseudobulk meta-analysis. The two resulting ranked gene lists were then intersected within each T cell gene list (CD4, CD8 separately), retaining only genes that (i) passed the significance threshold and (ii) showed the same direction across both.

Gene set enrichment analysis was performed on the full REML meta-analysis-ranked gene lists using fgsea (1.34.2). Genes were ranked by meta-LFC (NR vs R) and three curated gene set libraries were used: the MSigDB Hallmark collection [37], the Menyhart Core and Integrated cancer hallmark gene sets [38], and the ImmPort immunology gene set compendium [39]. Gene sets with fewer than 10 or more than 5,000 members were excluded; enrichment scores were estimated using 10,000 permutations. fgsea was run independently on the pseudobulk and single-cell REML-ranked lists for CD4 and CD8 T cells. Pathways were considered high-confidence if they reached nominal p < 0.05 with a consistent normalised enrichment score (NES) direction in both the pseudobulk and single-cell analyses.

### Dataset-level signature scoring

For queried gene signature, predefined patient-level pseudobulk expression matrices (details in supplementary methods S16) of single datasets were used to calculate signature enrichment scores. Expression matrices were organized with genes as rows and patients as columns. Signatures with fewer than three matched genes were excluded. Single-sample gene set enrichment analysis (ssGSEA) [40], implemented using the GSVA framework [41], was then applied to each patient to generate a continuous enrichment score for the corresponding gene signature. Differences between R and NR were assessed using the Wilcoxon rank-sum test. The magnitude of the difference was quantified using the mean score difference and Cohen’s (d) based on the pooled standard deviation.

### External validation cohort and preprocessing

Two independent bulk RNA-seq cohorts were used for external validation of the identified signature.

1. GSE223655 [42], comprised pre-treatment leukapheresis PBMC samples from 12 DLBCL patients treated with CAR-T cell therapy (6 complete responders [CR] and 6 non-responders [NR]; progressive disease relabelled NR for consistency). FPKM-normalized expression matrices and clinical metadata were obtained from GEO; expression values were log2(FPKM + 1)-transformed prior to analysis. For each patient, expression profiles across four sorted T cell subtypes (Naïve CD4, Naïve CD8, Non-Naïve CD4, Non-Naïve CD8) were averaged within subtype, yielding a single representative transcriptomic vector per patient per subtype, as well as an overall per-patient profile. ssGSEA scores were computed independently for the UP-in-NR and UP-in-R gene sets using the GSVA package (v2.2.1; ssgseaParam, normalize = TRUE), and a combined signature score was defined as UP_ssGSEA − DN_ssGSEA, where higher values indicate a more NR-like transcriptional state.
2. GSE325315 [43], comprised pre-treatment cell-free RNA (cfRNA) bulk RNA-seq profiles from 91 patients (64 R, 27 NR). Raw counts were TMM-normalised and log-transformed to counts per million (logCPM) using edgeR, with a prior count of 1. Ensembl gene IDs were mapped to HGNC symbols and duplicate mappings were resolved by retaining the entry with the highest mean expression. ssGSEA was run across all samples simultaneously, and combined scores were computed as above. Group differences in combined signature score were assessed using the Wilcoxon rank-sum test and effect size was quantified using Cohen’s d.

Further, two experimental datasets were included to evaluate the signature across CAR-T exhaustion and genetically perturbed in-vitro CAR-T cells.

3. GSE160154 [44] profiled CD8+ CAR-T cells bulk RNA-seq from 4 donors at Day 0 (pre-exhaustion control, n=8 samples) and Day 28 after chronic stimulation (exhausted, CS CAR-T+ and IS CAR-T+, n=15 samples); adjusted count matrices were log2(counts+1)-transformed.
4. GSE147991 [45] comprised CD19 CAR-T bulk RNA-seq from 3 donors across two conditions: Ctrl gRNA CART19 (n=3, functionally exhausted/NR-like) and GM-CSF KO CART19 (n=3, enhanced function/R-like). For both cohorts, ssGSEA was run across all samples simultaneously using the same parameters described above.

### Database architecture and web interface implementation

ImmunoCAR Atlas is built on an Nginx server (version 1.18) installed on a machine running Ubuntu 22.04.5 LTS (Jammy Jellyfish) with Linux as the operating system. The responsive front-end, suitable for mobile, tablet, and desktop platforms, was developed using HTML5, CSS3, and JavaScript. The back-end of the platform is controlled using Flask (version 3.0.3), which manages server-side logic, routing, and data processing. Flask integrates the front-end templates with the back-end functionalities, enabling dynamic rendering of content and seamless interaction with the database. Data for the platform’s 16 reference atlases including expression matrices (AnnData/.h5ad), differential expression tables, cell-composition and pathway enrichment results, and clinical metadata are stored as structured flat files rather than a conventional database, indexed by dataset, tissue, timepoint, and response group for fast retrieval. Interactive visualizations (UMAPs, heatmaps, DEG and pathway plots) are built with Plotly and Dash, while the Patient Cell Composition module uses R (tidyverse, ggplot2) called via rpy2. A Flask-based REST API allows programmatic querying of metadata, DEG/pathway results, and h5ad downloads across all 16 atlases.

### AI-assisted querying of Atlas

To enable natural-language querying of the ImmunoCAR Atlas, we implemented a Model Context Protocol (MCP) server: a lightweight, independent translation layer that exposes the database’s existing REST API as a defined set of callable tools. Deployed on the same host as the database and reverse-proxied through nginx at a dedicated public endpoint, the MCP server receives tool-call requests from a large language model (Claude, Anthropic), converts them into standard read-only HTTP requests against the local REST API, and returns structured JSON responses to the model. This allows any authorized user to query datasets, differential expression results, and other precomputed analyses through conversational prompts, without modifying, restarting, or otherwise altering the underlying application or database. Further details are provided in Supplementary Methods S17.

## Results

### The ImmunoCAR Atlas captures the clinical, cellular and therapeutic diversity of CAR-T single-cell studies

To capture the diversity and technical heterogeneity of published CAR-T single-cell studies, the ImmunoCAR Atlas assembles 43 scRNA-Seq datasets spanning 5.7 million cells from 584 patients treated with 12 CAR-T cell therapies including tisagenlecleucel (tisa-cel), axicabtagene ciloleucel (axi-cel), ciltacabtagene autoleucel (cilta-cel), and idecabtagene vicleucel (ide-cel) across diffuse large B-cell lymphoma (DLBCL), multiple myeloma (MM), and acute lymphoblastic leukemia (ALL) (Figure 1A). Critically, the compendium covers the full clinical trajectory: pre-treatment baseline, manufactured infusion product, and post-treatment follow-up, sourced from peripheral blood, bone marrow, and cell products. This breadth positions the atlas as a resource not merely for aggregation, but for systematic, cross-study investigation of therapy-associated immune remodeling over time.

**Figure 1.**
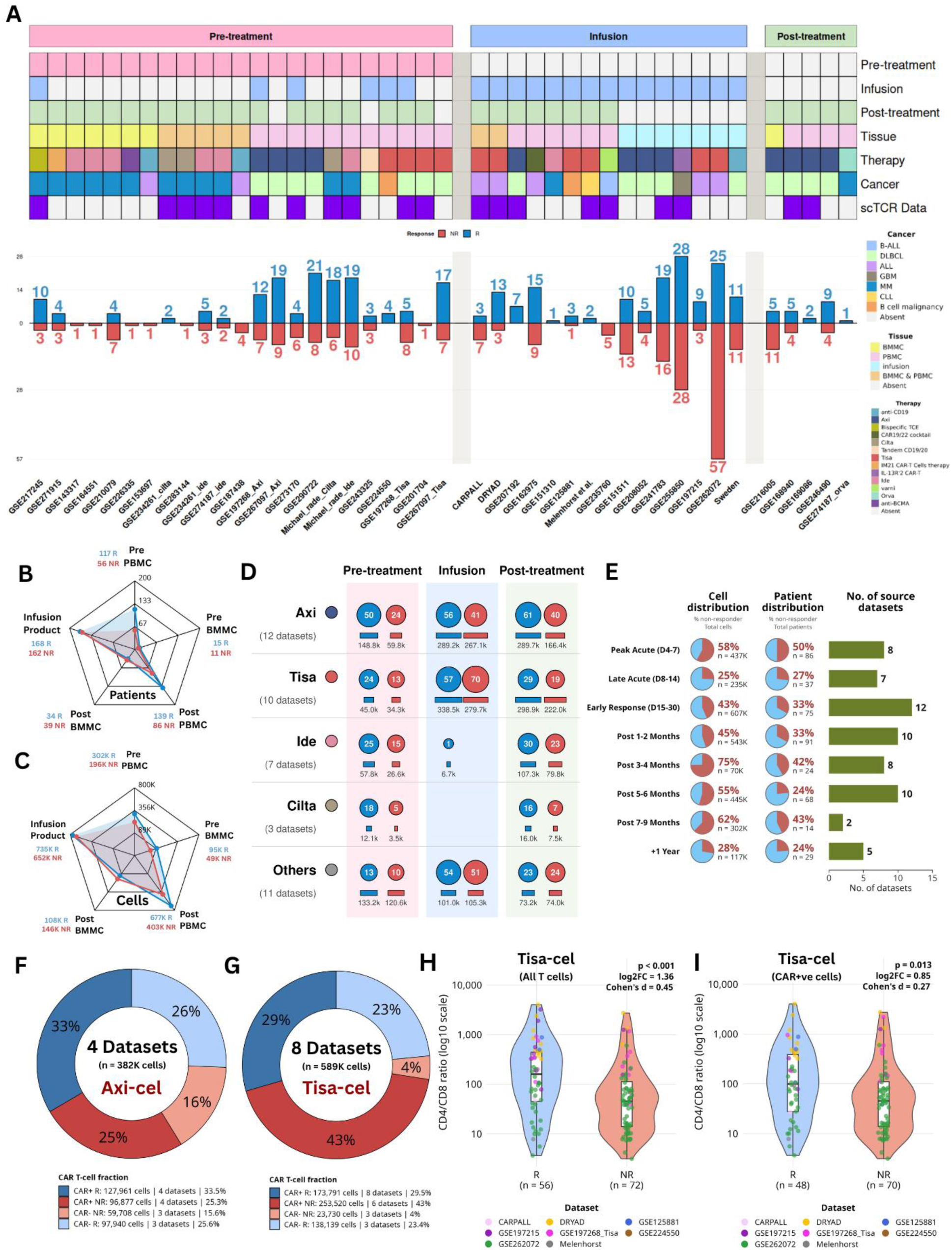
**(A).** Overview of the single-cell RNA sequencing (scRNA-seq) compendium analyzed in this study. Each column represents an individual dataset. (**B,C).** Radar plot depicting distribution of patients (B) and cells (C) across sampling compartments, stratified by clinical response. (**D).** Patient and cell representation across treatment phases, stratified by CAR-T product. Circles indicate patient numbers, with blue and red denoting R and NR, respectively; bars below indicate corresponding cell numbers. (**E).** Longitudinal granular distribution of post-infusion sampling timepoints across the compendium. Pie charts show the proportion of R/NR cells and patients, while horizontal bars indicate the number of contributing datasets. (**F,G)** CAR-positive and CAR-negative T-cell fractions in axi-cel and tisa-cel infusion products, stratified by response. (**H,I)** CD4/CD8 ratios in tisa-cel-derived all T cells and CAR-positive T cells, respectively, comparing R and NR groups.

Clinically, compendium reveals substantial variation in the clinical and sampling architecture of published CAR-T studies. At the patient level (Figure 1B), peripheral blood provides the largest representation at both baseline (117 responders (R) and 56 non-responders (NR)) and follow-up (139 R and 86 NR), whereas the infusion-product compartment includes 168 R and 162 NR. Bone marrow sampling provides an additional dimension, with 15 R and 11 NR represented before treatment and 34 R and 39 NR after treatment. At the cellular level, these compartments encompass markedly different numbers of profiled cells, with the infusion product alone contributing approximately 1.39 million response-annotated cells (Figure 1C), highlighting the distinction between patient and sample representation.

The architectural design of the atlas emphasizes longitudinal resolution: five datasets include all three treatment phases, while another 15 contain paired pre- and post-treatment samples, of which 13 represent both R and NR populations (Supplementary Fig. S1A). Longitudinal coverage also varies across therapeutic modalities, with axi-cel and tisa-cel having the broadest representation across treatment phases (Figure 1D). For example, axi-cel includes 56 R and 41 NR in the infusion-product compartment, whereas the corresponding post-treatment collections extend this representation to 61 R and 40 NR. This modality-resolved structure provides a basis for distinguishing shared CAR-T-associated immune features from product-specific cellular characteristics, while retaining the underlying study and sampling context.

Beyond transcriptomic breadth, the atlas is architectured for mechanistic depth. Twenty datasets include matched single-cell TCR sequencing (scTCR-seq), enabling T cell clonotype tracking and clonal expansion analyses (Figure 1A). Furthermore, data were generated across seven sequencing platforms where Illumina NovaSeq accounted for 62.7% of total datasets and six library preparation methods. These technical variables are catalogued as essential metadata, ensuring that integrated analyses can appropriately account for batch and platform effects. Demographic metadata are available for a substantial subset of patients: age data (Supplementary Figure S1B) for 54.6% (n = 319, median 51 years) and gender data (Supplementary Figure S1C) for 40.6% (n = 237, 59% male).

A particularly important feature of the atlas is its coverage of clinically distinct post-infusion intervals (Figure 1E). Sampling extends from acute monitoring (days 4–14) and early response assessment (days 15–30) to follow-up beyond one year. Early response assessment is the most extensively represented interval, comprising approximately 607K cells from 75 patients across 12 datasets. Notably, the distribution of non-responder-derived cells varies substantially across intervals, ranging from 25% during late acute monitoring to 75% at 3–4 months. Patient-level representation shows a corresponding, but not identical, pattern, with non-responders accounting for 24–50% of patients across the sampled intervals. This distinction between cell-level and patient-level representation is important: cell abundance does not necessarily translate into equivalent clinical representation, underscoring the need for patient-aware statistical approaches in cross-study analyses. Nevertheless, the availability of samples extending beyond one year provides an opportunity to investigate immune states associated with both early post-infusion dynamics and longer-term clinical outcomes.

The atlas additionally enables interrogation of the manufactured therapeutic product. Four axi-cel datasets contribute approximately 382,000 T cells, while eight tisa-cel datasets contribute approximately 589,000 T cells (Figure 1F,G). CAR expression is heterogeneous within these products: CAR-positive cells constitute 58.8% of the analysed axi-cel T cells and 72.5% of tisa-cel T cells, with the remaining fractions comprising CAR-negative cells. These distributions expose a substantial CAR-negative fraction within the manufactured products and demonstrate that the cellular composition of an infused product cannot be adequately represented by treating all constituent T cells as a homogeneous CAR-expressing population. The differing proportions of CAR-positive and CAR-negative cells across products further motivate analyses that explicitly distinguish these compartments when investigating therapeutic cellular phenotypes. Consistent with the importance of product composition, response-associated differences were also evident in the CD4/CD8 balance of tisa-cel-derived T cells (Figure 1H,I). Across all T cells, responders exhibited a significantly higher CD4/CD8 ratio than non-responders (P < 0.001; log₂ fold change = 1.36; Cohen’s d = 0.45). This association remained detectable when the analysis was restricted to CAR-positive T cells (P = 0.013; log₂ fold change = 0.85; Cohen’s d = 0.27), although the effect size was smaller. Collectively, the ImmunoCAR Atlas integrates the clinical, temporal and cellular dimensions of CAR-T single-cell studies within a common analytical framework. By connecting pre-treatment immune states, infusion-product composition and longitudinal post-infusion profiles with patient-level response annotations, it provides a foundation for investigating how baseline immune context, therapeutic product heterogeneity and post-infusion immune remodelling intersect to shape CAR-T treatment outcomes.

### Architecture of ImmunoCAR Atlas

ImmunoCAR Atlas was developed as a modular, patient-stratified single-cell resource integrating harmonized transcriptomic data with curated clinical metadata and CAR-T treatment outcomes (Figure 2). Its architecture comprises five interconnected components: (i) data collection and curation; (ii) standardized processing and annotation to harmonize cell identities, T-cell states, and clinical response categories across studies; (iii) analytical modules operating at dataset, cross-cohort, and atlas-wide levels; (iv) query and evaluation tools for gene- and signature-level exploration; and (v) data access through interactive browsing, search, and download interfaces, complemented by a programmatic API and large language model (LLM)-enabled natural-language querying.

**Figure 2.**
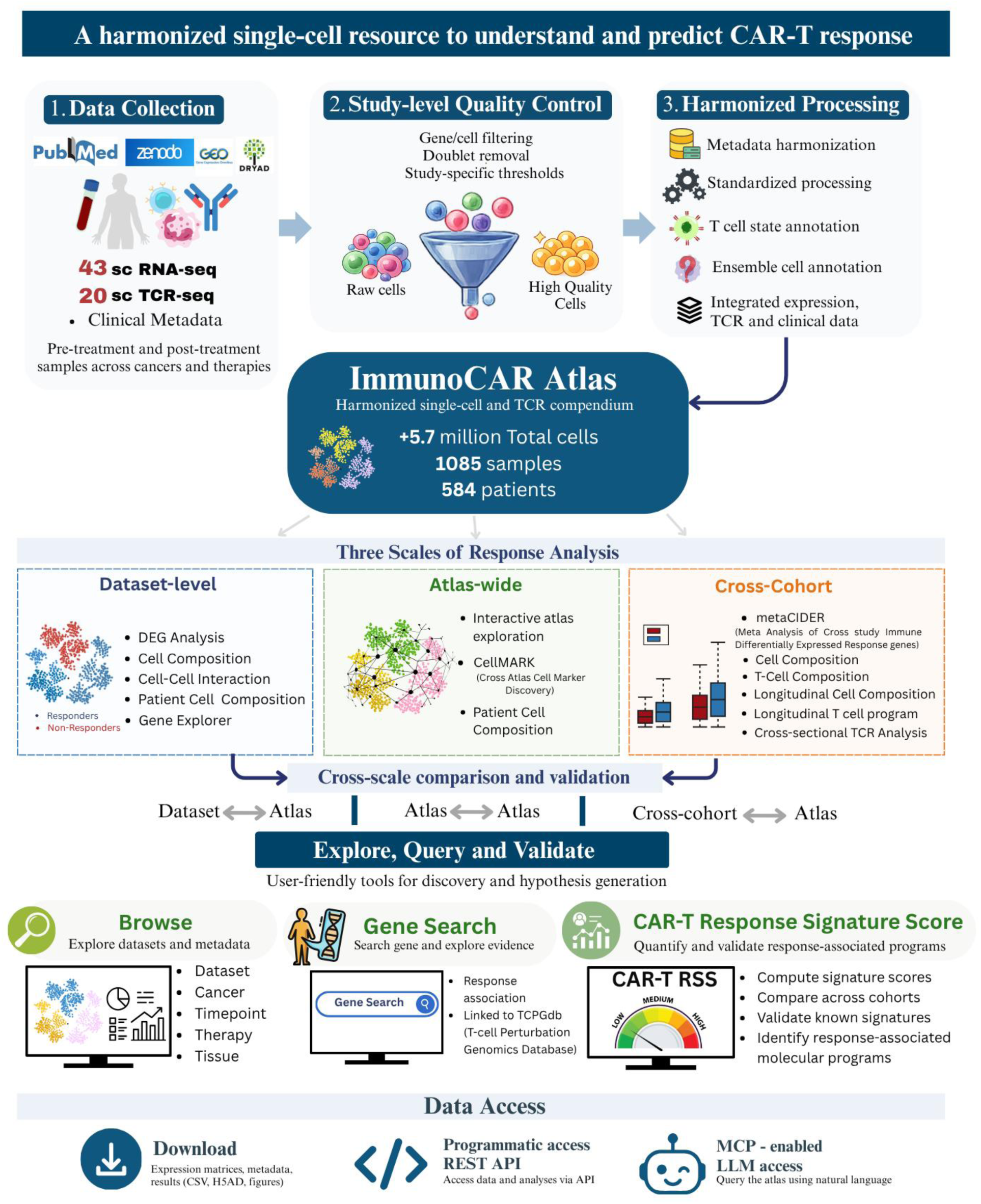
Architecture of the ImmunoCAR Atlas, integrating data curation, standardized processing and annotation, multi-tier analytics, and interactive data access.

The analytical framework is organized into three complementary tiers (Figure 2). Dataset-level analysis enables responder-versus-non-responder (R vs NR) comparisons within individual cohorts, including differential gene expression (DEG), cell composition profiling, infer cell–cell communication networks, patient-level immune populations assessment, and query gene expression across cell states and response groups via an interactive Gene Explorer. Cross-cohort analysis extends these investigations across studies sharing comparable clinical or biological variables. This includes overall and T-cell-specific composition profiling, longitudinal tracking of immune populations and T-cell programs across treatment timepoints, and cross-sectional T-cell receptor (TCR) analysis to investigate clonal expansions associated with clinical response. At this level, metaCIDER (Meta-analysis of Cross-study Immune Differentially Expressed Response genes) aggregates dataset-level differential expression results to identify genes reproducibly associated with response or resistance across independent cohorts. The resulting gene sets can subsequently be evaluated using the custom signature scoring module to assess their consistency and predictive potential across datasets. Atlas-wide analysis leverages 16 scVI-integrated reference atlases to increase the number of patients and cells available for identifying response-associated immune features, thereby improving the ability to detect signals that may be underpowered in individual cohorts. This tier includes patient- and cell-type-level composition analyses to identify immune populations reproducibly enriched in responders or non-responders, complemented by CellMARK (Cross-Atlas Cell Marker Discovery) for identifying response-associated cell-type markers across integrated atlases. A distinguishing feature of the database is a gene-centric search interface provides a consolidated report card for any queried gene, summarizing its prevalence and direction of differential expression across datasets and comparisons.

Beyond interactive analysis, ImmunoCAR Atlas provides downloadable, versioned, analysis-ready integrated reference objects. These can be directly incorporated into external single-cell workflows for reference mapping, label transfer, and downstream computational modelling. Together, these complementary analytical and access components provide a framework for investigating response-associated immune features at individual-cohort and cross-study scales, while facilitating independent validation and downstream translational applications.

### Constructing atlases across diverse therapies, cancer types, tissues, and timepoints

To enable systematic integration of immune landscapes across CAR-T responder and non-responder datasets, we developed a standardized four-stage pipeline for dataset curation, metadata harmonization, cell annotation, and scVI integration (Figure 3A). Applying this framework, we generated 16 cancer-, tissue-, and timepoint-specific reference atlases spanning Axi-cel in DLBCL, Tisa-cel in ALL, and ide-cel/cilta-cel in multiple myeloma (Figure 3B). Atlas size ranged from ∼67K to >830K cells, and the proportion of responder-derived cells varied widely across atlases, reflecting the natural class imbalance present in the underlying published cohorts and underscoring the importance of integration to standardize datasets.

**Figure 3.**
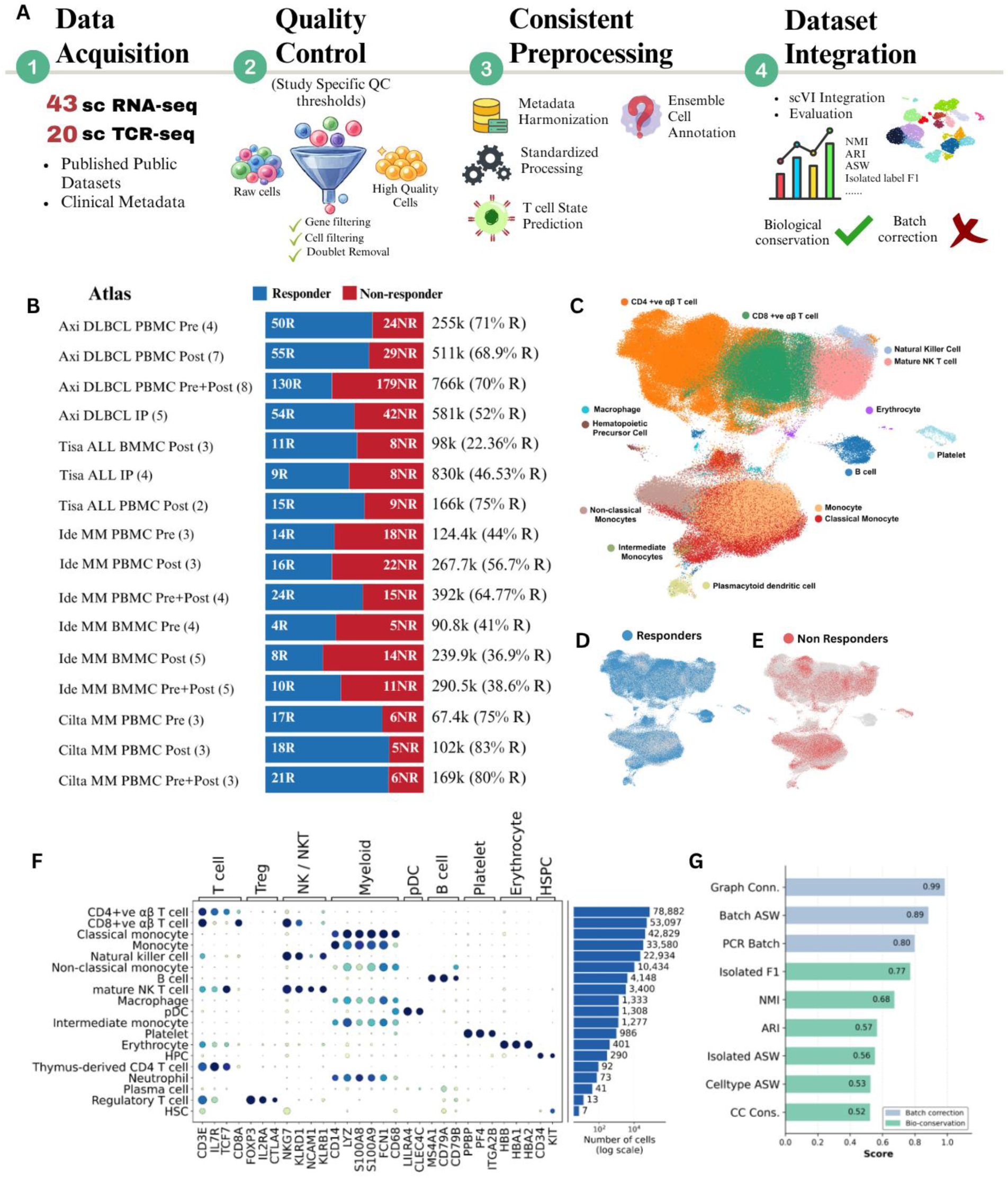
**A)** Dataset curation, harmonization, annotation, and scVI integration workflow. **B)** Overview of 16 CAR-T atlases across cancer, tissue, and timepoint strata. **C)** Representative Axi-cel DLBCL PBMC pre-infusion atlas, showing cell types and clinical outcomes. **D–E)** Responder and non-responder UMAPs. **F)** Canonical marker validation of cell-type annotations. **G)** scIB metrics assessing integration quality.

As a representative example, the Axi-cel DLBCL PBMC pre-infusion atlas, integrating four studies, resolved major immune populations including T cells, NK cells, myeloid cells, B cells, dendritic cells, and progenitor populations (Figure 3C). Upon splitting cells by clinical outcome, responder-(Figure 3D) and non-responder-derived cells (Figure 3E) were broadly intermixed within clusters, indicating effective correction of study- and patient-level batch effects while preserving biological structure. Cell-type annotations were supported by canonical lineage markers (Figure 3F), with expected marker expression for T cells (*CD3E, IL7R, TCF7*), NK cells (*NKG7, KLRD1*), myeloid cells (*CD14, LYZ*), B cells (*MS4A1, CD79A*), pDCs (*LILRA4, CLEC4C*), and HSPCs (*CD34, KIT*). Marker expression was largely restricted to the corresponding populations, supporting annotation accuracy across both abundant and rare cell types.

Integration quality was assessed using scIB metrics [30] (Figure 3G). High batch-correction scores, including graph connectivity (0.99), batch ASW (0.89), and PCR batch (0.80), indicated effective reduction of technical variation, while moderate-to-high biological-conservation scores, including isolated-label F1 (0.77), NMI (0.68), and ARI (0.57), confirmed preservation of cell identity and biological structure. Together, these results demonstrate robust integration across studies while retaining biologically meaningful cell states, providing a consistent foundation for downstream response analyses. Integration metrics for all atlases are available through the atlas dashboard at https://car-t-atlas.igib.res.in/atlas/

### Cross-Atlas Discovery of Robust Cell-Type Markers with CellMARK

An open challenge in CAR-T research is distinguishing reproducible cell-type marker signals from dataset-specific noise. Atlas-level composition analysis addresses this by testing cell-type associations with response/resistance across an integrated object built from many cohorts, surfacing effects that individual studies are often underpowered to detect on their own.

We used CellMARK to investigate baseline (pre-infusion) PBMC composition across three clinically relevant settings: axi-cel-treated DLBCL, tisa-cel-treated DLBCL, and ide-cel-treated multiple myeloma (MM) patients (Figure 4A–C, respectively). Cross-atlas analysis revealed both shared and therapy- or cancer-specific patterns of response-associated cellular composition. In DLBCL, responders receiving either axi-cel or tisa-cel exhibited higher baseline abundances of B cells and non-classical monocytes (Figure 4A–B), suggesting that these populations may represent shared cellular correlates of favourable CAR-T outcomes across CD19-directed therapies. Natural killer (NK) cells also showed a responder-associated enrichment in the axi-cel atlas, with a similar, albeit less pronounced, trend in the tisa-cel atlas. Conversely, the tisa-cel atlas highlighted greater baseline representation of macrophages and classical monocytes in non-responders, suggesting a potentially unfavourable myeloid-associated immune context. Extending this analysis to BCMA-directed CAR-T therapy in MM, the ide-cel atlas revealed higher baseline CD8+ T-cell abundance in responders, whereas macrophages were more prominently represented in non-responders (Figure 4C). Together, these observations highlight the ability of CellMARK to capture both potentially shared response-associated immune features and context-dependent cellular differences across CAR-T therapies and cancer types.

**Figure 4.**
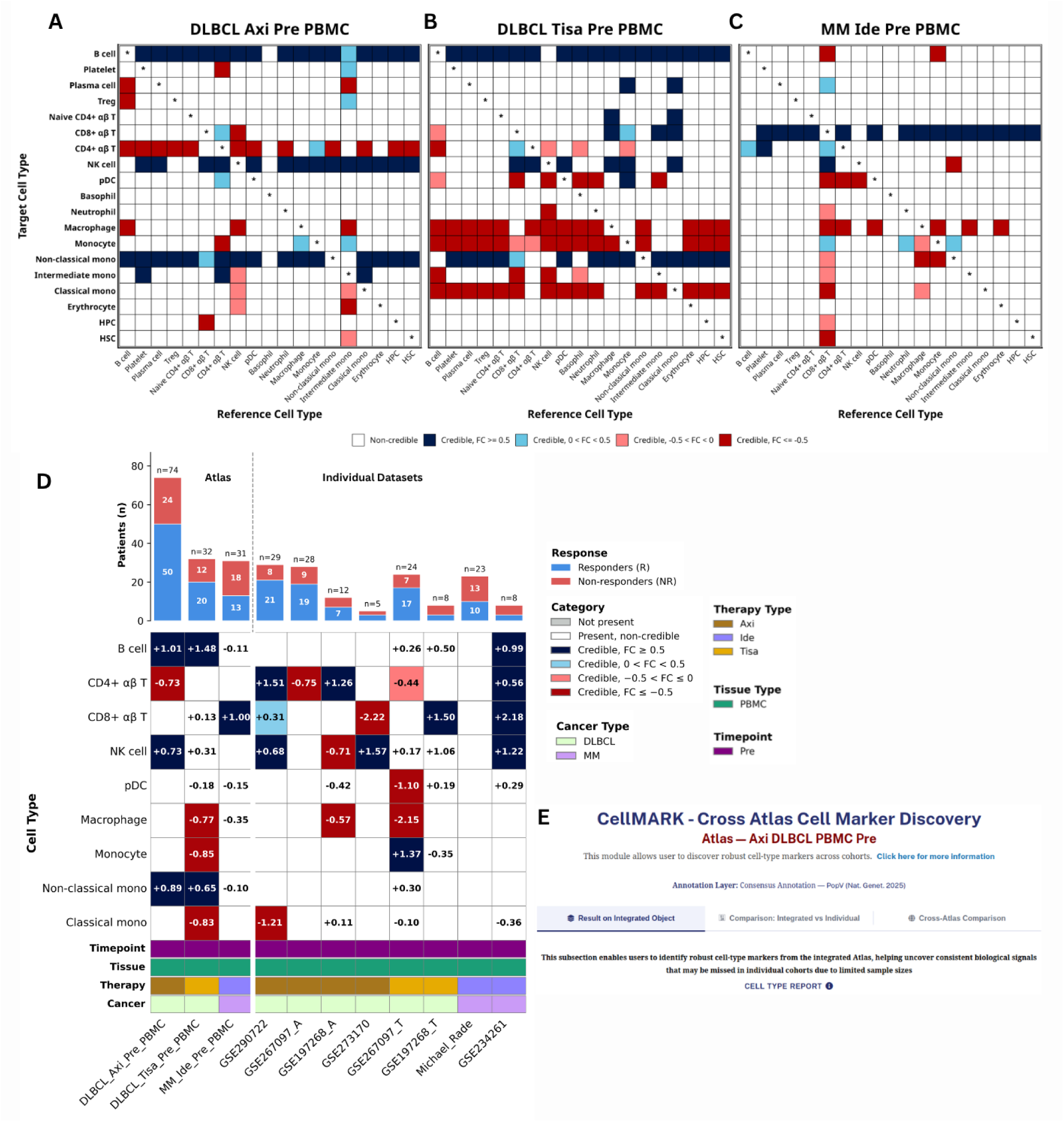
**(A-C)** Credibility heatmaps summarizing cell-type compositional associations with CAR-T response within three integrated pre-infusion PBMC atlases: (A) axi-cel-treated DLBCL, (B) tisa-cel-treated DLBCL, and (C) ide-cel-treated multiple myeloma (MM). Heatmaps illustrate the direction and credibility of response-associated cell-type enrichment across references (each cell type). **(D)** Comparison of compositional analysis results obtained from individual cohorts versus their corresponding integrated atlases **(E)** CellMARK user interface for the Axi DLBCL PBMC Pre atlas.

Importantly, the added value of CellMARK became apparent when these integrated atlas-level findings were compared with compositional analyses performed independently in the constituent datasets (Figure 4D). Several associations that were weak or statistically inconclusive in individual cohorts gained stronger statistical support following integration. For example, non-classical monocytes did not consistently meet the predefined credibility (FDR < 0.05) in the individual axi-cel and tisa-cel cohorts; however, following integration, this population emerged as a statistically supported responder-associated feature in both atlases. This illustrates how cross-cohort integration can overcome limitations imposed by individual cohort sizes, improving the ability to detect cellular associations that might otherwise remain obscured.

Collectively, these results demonstrate that CellMARK (Figure 4E) provides more than a consolidated view of immune-cell composition: it offers a systematic framework for uncovering and prioritizing response-associated cellular populations across studies. By distinguishing signals supported across integrated cohorts from those that are more context-specific, CellMARK facilitates the identification of candidate cellular biomarkers that warrant further investigation and independent validation in CAR-T therapy.

### metaCIDER Identifies a Reproducible Pre-Infusion Response Signature in anti CD19–Treated DLBCL

Identifying genes and molecular correlates reproducibly associated with CAR-T response and resistance is essential for understanding the biological determinants of treatment outcomes and developing robust predictive biomarkers. However, differences in cohort composition, clinical characteristics, cell-type annotation, and differential-expression methodologies often complicate the identification of consistent response-associated signals across independent studies. To address this challenge, we developed metaCIDER (Meta-analysis of Cross-study Immune Differentially Expressed Response genes), a standardized framework that harmonizes study selection and differential-expression criteria to identify genes exhibiting consistent response-associated regulation across independent cohorts (Figure 5A).

**Figure 5.**
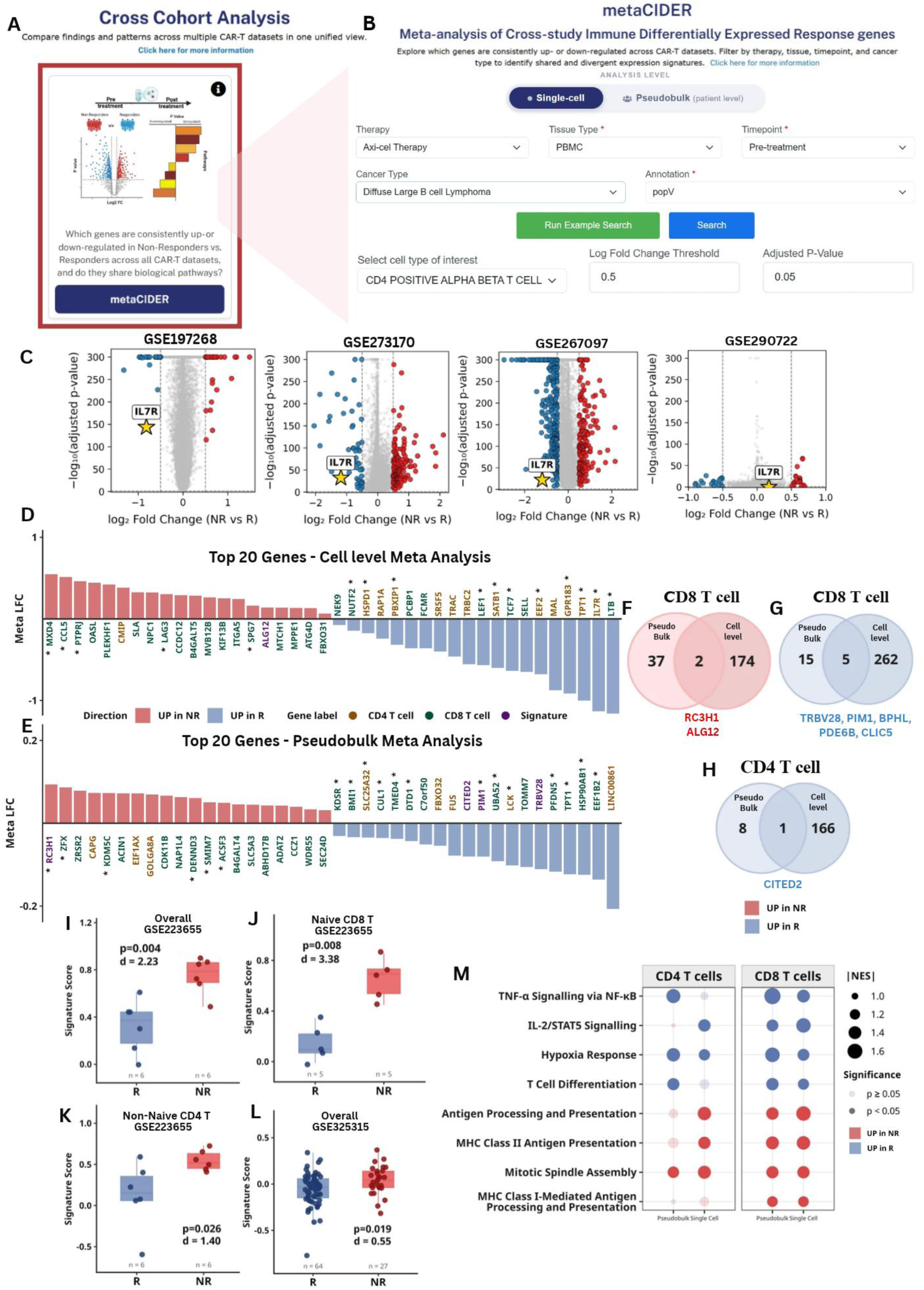
**A)** Accessing the metaCIDER module within the Cross-Cohort Analysis. **B)** metaCIDER query interface showing filters for therapy, tissue, timepoint, and cancer. **C)** Representative volcano plot highlighting IL7R, significantly and concordantly upregulated in responders (R) across datasets. **(D)** Bar plot of the top 20 genes ranked by combined score (−log₁₀ p × |meta-LFC|) from the cell level meta-analysis across six pre-treatment DLBCL PBMC cohorts, shown for CD4 and CD8 T cells combined. Bars are coloured by direction (UP in NR, red; UP in R, blue); gene labels are coloured by cell-type origin (amber = CD4, teal = CD8, purple = signature genes); asterisk (*) denotes TPS score > 0.1 in CD8T, CD4T, or CAR-T. **(E)** As in (D), for the pseudobulk REML meta-analysis. **(F-G)** Venn diagram showing overlap of NR/R genes between pseudobulk and single-cell CD8 T cell meta-analyses. **(H)** in CD4 T cells **(I)** Combined ssGSEA signature score (UP − DN) in the independent bulk cohort GSE223655 (n = 12; 6 CR, 6 NR) across all T cells. Wilcoxon p = 0.004, Cohen’s d = 2.23. **(J)** Signature score stratified by Naive CD8 T cells in GSE223655. **(K)** Signature score stratified by Non-Naive CD4 T cells in GSE223655. **(L)** Validation in pre-treatment cfRNA cohort GSE325315. **(M)** Dot plot of pathways identified by fgsea using pseudobulk and single-cell REML-ranked gene lists. Dot size encodes |NES|; opacity reflects nominal significance (opaque = p < 0.05); colour indicates direction (red = UP in NR, blue = UP in R).

We first evaluated metaCIDER in a focused analysis of axi-cel–treated DLBCL, restricting the query to pre-infusion PBMC-derived CD8+ αβ T cells (Figure 5B). Applying the default thresholds of |log₂FC| > 0.5 and adjusted *P* < 0.05, four independent cohorts met the eligibility criteria for cross-study analysis. Despite differences in the number and magnitude of differentially expressed genes across individual cohorts, metaCIDER identified reproducible response-associated signals. For example, *IL7R* was significantly upregulated in responders in three of the four cohorts (75%), illustrating the ability of the framework to distinguish recurring biological signals from cohort-specific findings (Figure 5C).

Having established its ability to recover reproducible signals within a single CAR-T product, we next expanded the analysis across axi-cel and tisa-cel to identify transcriptional features associated with response to anti-CD19 CAR-T therapy more broadly. Across six independent pre-infusion DLBCL datasets, we examined CD4 and CD8 T-cell compartments at both single-cell and pseudobulk resolutions. This analysis revealed several genes such as *GPR183* [46], *TCF7* [47], and *LEF1* [48] emerged among the top ranked genes and exhibited T-cell perturbation scores (TPS) >0.1 in at least one of the CD4, CD8, or CAR-T cell contexts [49], highlighting candidates with potential functional relevance to T-cell biology. The top 20 genes from the cell level and pseudobulk level meta-analyses are presented in Figures 5D and 5E, respectively.

To identify the most robust candidates, we focused on genes showing concordant response-associated expression across both single-cell and pseudobulk analyses. In CD8 T cells, two genes associated with non-response, *RC3H1* and *ALG12* (Figure 5F). Conversely, five genes are associated with response *TRBV28*, *PIM1*, *BPHL*, *PDE6B*, and *CLIC5* (Figure 5G). Within CD4 T cells, *CITED2* emerged as the sole gene concordantly upregulated in responders (Figure 5H). Together, these eight genes constituted a cross-resolution candidate signature, with distinct transcriptional features associated with subsequent response and resistance.

We next investigated whether this signature retained its discriminatory potential in independent patient cohorts and across transcriptomic platforms. In the independent GSE223655 bulk RNA-seq apheresis cohort (n = 12; 6 R and 6 NR), the signature significantly distinguished response groups at the overall (p = 0.004, Cohen’s d = 2.23; Figure 5I). The separation was particularly pronounced in naive CD8 T cells (p = 0.008, Cohen’s d = 3.38; Figure 5J) and non-naive CD4 T cells (p = 0.026, Cohen’s d = 1.40; Figure 5K). Directional consistency was also observed in naive CD4 and non-naive CD8 T-cell populations (Supplementary Figures 2A–B). Importantly, the signature also retained significance in the independent pre-treatment cell-free RNA (cfRNA) cohort GSE325315 (n = 91; 64 R and 27 NR; p = 0.019, Cohen’s d = 0.55; Figure 5L). The recovery of the response-associated signal in both cellular transcriptomes and circulating RNA provides evidence of its generalizability across distinct sample types and transcriptomic platforms.

To further explore the biological relevance of the non-responder-associated transcriptional state, we examined the signature in two independent CAR-T perturbation datasets. In GSE160154, exhausted CAR-T cells exhibited a higher combined non-responder signature score than control cells (Cohen’s d = 0.28; Supplementary Figure 2C), consistent with the expected direction of an exhaustion-associated transcriptional state. In GSE147991, control gRNA cells exhibited higher signature scores than GM-CSF knockout cells (Cohen’s d = −0.82; Supplementary Figure 2D), consistent with attenuation of the non-responder-associated transcriptional state following GM-CSF knockout. Neither comparison reached statistical significance (p>0.05), consistent with the small sample sizes and in vitro context.

Finally, we investigated the broader biological programs associated with response and resistance (Figure 5M). Responder-associated programs included TNF-α/NF-κB and IL-2/STAT5 signalling, hypoxia response, and T-cell differentiation. In contrast, non-responder-associated programs included antigen processing and presentation, MHC class I and II pathways, and mitotic spindle assembly. These pathway patterns were broadly concordant across CD4 and CD8 T-cell compartments, suggesting that the transcriptional differences associated with subsequent CAR-T outcomes extend beyond individual genes to encompass coordinated immune and cellular programs.

## DISCUSSION

The single-cell CAR-T literature has grown rapidly but unevenly: dozens of studies, most too small to power outcome analyses on their own, scattered across inconsistent metadata, tissue sources, sampling timepoints, and annotation schemes [50]. Consequently, it remains unclear which reported associations represent reproducible features of response and which are specific to individual cohorts. ImmunoCAR Atlas addresses this challenge by bringing together a harmonized, clinically annotated collection of CAR-T single-cell datasets and providing analytical frameworks to interrogate response-associated features across independent studies. Its value lies not only in consolidating these datasets, but also in enabling candidate biomarkers to be evaluated in the context of the broader CAR-T landscape.

A central insight emerging from the CellMARK analysis is that the immune composition of patients before CAR-T infusion may provide information about subsequent therapeutic outcomes. The convergence of B-cell and non-classical monocyte enrichment in responders across anti-CD19-treated DLBCL cohorts points to potentially shared features of a favourable pre-infusion immune environment across different CAR-T products. The additional enrichment of NK cells in axi-cel responders, together with the distinct myeloid associations observed in tisa-cel and BCMA-directed multiple myeloma cohorts, suggests that cellular correlates of response may also depend on the therapeutic and disease context. These observations argue against interpreting CAR-T response through a single universal immune-composition signature. Rather, they highlight the importance of distinguishing cross-cohort associations from those that are specific to a particular treatment setting. Importantly, the associations identified through integrated analysis warrant further investigation to establish their functional relevance and predictive value.

Cellular composition, however, captures only one dimension of the pre-treatment immune landscape. The metaCIDER analysis addresses a complementary question: whether transcriptional features associated with response and resistance can be recovered across independent cohorts and analytical resolutions. The identification of a concordant eight-gene signature across pre-infusion anti-CD19 DLBCL datasets, followed by its evaluation in independent bulk RNA-seq and cell-free RNA-seq datasets, provides an additional level of evidence for the reproducibility of these associations. The recovery of related signals across single-cell and bulk measurements is particularly encouraging because it suggests that clinically relevant transcriptional features need not be restricted to single-cell profiling. Nevertheless, cross-platform validation should be distinguished from prospective clinical validation, and the diagnostic and predictive utility of this signature will require evaluation in larger, independently assembled patient cohorts.

The biological context of the signature further suggests that response and resistance may be associated with different pre-existing T-cell states before CAR-T manufacture. Among the candidate genes, PIM1 is of particular interest given its established involvement in the NF-κB–PIM1–Eomesodermin axis that supports CD8⁺ T-cell memory maintenance and function [51]. CITED2, a stemness-associated transcription factor upregulated in memory and stem-like CAR-T cell subsets, contributes to transcriptional programmes associated with quiescence and persistence [52]. RC3H1 encoding a post-transcriptional repressor whose combined disruption with Regnase-1 enhances CAR-T antitumour activity and persistence in engineered human T cells [53]. Its upregulation in NR pre-treatment CD8 T cells may suggest predisposition to suppression before CAR-T therapy. However, the functional consequences of these associations cannot be inferred from baseline expression alone. Determining whether these genes actively influence T-cell fitness, persistence or susceptibility to dysfunction will require targeted functional studies.

The accompanying pathway-level analyses provide a broader perspective on these transcriptional differences. The enrichment of TNF-α signalling via NF-κB, IL-2/STAT5 signalling and hypoxia-associated programmes in responder-derived CD4⁺ and CD8⁺ T cells is consistent with the retention of transcriptional programmes involved in T-cell survival, cytokine responsiveness [54], differentiation plasticity [55] and metabolic adaptation [56]. These observations are relevant to CAR-T biology, where the differentiation state and metabolic fitness of the starting T-cell population can influence the properties of the manufactured product. Conversely, the enrichment of mitotic spindle assembly and antigen-processing and presentation programmes in non-responder-derived T cells suggests a distinct transcriptional state associated with subsequent treatment failure. Persistent antigen exposure and inflammatory signalling are established drivers of T-cell dysfunction and exhaustion [57]. However, these findings warrant further investigation through protein-level and functional analyses.

Taken together, the two case studies show two distinct but complementary uses of the atlas: cell-composition discovery and cross-cohort differential-expression discovery with cross-atlas and cross-platform validation both converging on candidate biomarkers that would have been difficult to nominate, and impossible to stress-test, from any single published dataset. This is the core value proposition of ImmunoCAR Atlas relative to prior resources: it is not only a larger catalogue of cells, but a framework in which every candidate association can be immediately checked for reproducibility before it is proposed as a biomarker.

Looking forward, we anticipate ImmunoCAR Atlas will grow along three major axes:: (i) broader disease and construct coverage; (ii) integration of single-cell multi-omic data to link epigenetic states, surface protein expression, and spatial architecture to response; (iii) establishment of a community contribution portal enabling researchers to deposit and share pre-publication datasets with standardized metadata.

In summary, by pairing a harmonized, pan-immune, outcome-linked cell atlas with built-in cross-cohort validation tools such as CellMARK, metaCIDER and CAR-T RSS, ImmunoCAR Atlas is intended to function less as a static reference and more as a discovery platform, where any candidate biomarker can be interrogated for reproducibility across the CAR-T field before it is carried forward to experimental or clinical follow-up.

### DATA AVAILABILITY

ImmunoCAR Atlas is freely accessible at https://car-t-atlas.igib.res.in/. All processed data, metadata tables, analytical module outputs, and scVI-integrated reference atlases are available for download through the web interface or programmatically via the REST API.

## ACKNOWLEDGEMENTS

The ImmunoCAR atlas was constructed on the basis of integrating public CAR-T therapy scRNA-seq data sets. We acknowledge all the researchers and participants to share their data sets. Sherry Bhalla acknowledges support for MMP075204 from the Council of Scientific and Industrial Research (CSIR). Kirti Jain acknowledges CSIR for fellowship support. Simran Jit acknowledges the University Grants Commission (UGC) for fellowship support. Tanishqua Neeraj Sharma acknowledges support from the CSIR-IGIB project GAP0277. Diksha Singh Rathore and Anas acknowledge support from the CSIR project MMP075204. Radul acknowledges CSIR-IGIB for support. Kirti Jain, Simran Jit and Sherry Bhalla gratefully acknowledge the Academy of Scientific and Innovative Research (AcSIR), Ghaziabad, India for academic support. We acknowledge the support from CSIR-IGIB for infrastructure and its IT department for computing facilities.

## AUTHOR’S CONTRIBUTIONS

Data Curation: Sherry Bhalla, Kirti Jain, Tanishqua Neeraj Sharma, Diksha Singh Rathore, Anas Ansari; Methodology: Sherry Bhalla, Kirti Jain, Anas Ansari, Simran Jit, Diksha Singh Rathore; Formal Analysis: Sherry Bhalla, Kirti Jain, Anas Ansari, Simran Jit, Diksha Singh Rathore, Tanishqua Neeraj Sharma; Visualization: Sherry Bhalla, Kirti Jain, Tanishqua Neeraj Sharma, Diksha Singh Rathore, Radul R. Dev; Writing: Sherry Bhalla, Kirti Jain, Tanishqua Neeraj Sharma, Anas Ansari, Simran Jit, Diksha Singh Rathore; Review & Editing: Sherry Bhalla, Kirti Jain, Diksha Singh Rathore; Resources & Funding acquisition: Sherry Bhalla.

## SUPPLEMENTARY DATA

1. Supplementary Figure S1

2. Supplementary Figure S2

3. Supplementary Methods

## Supplementary Figures

**Figure S1.**
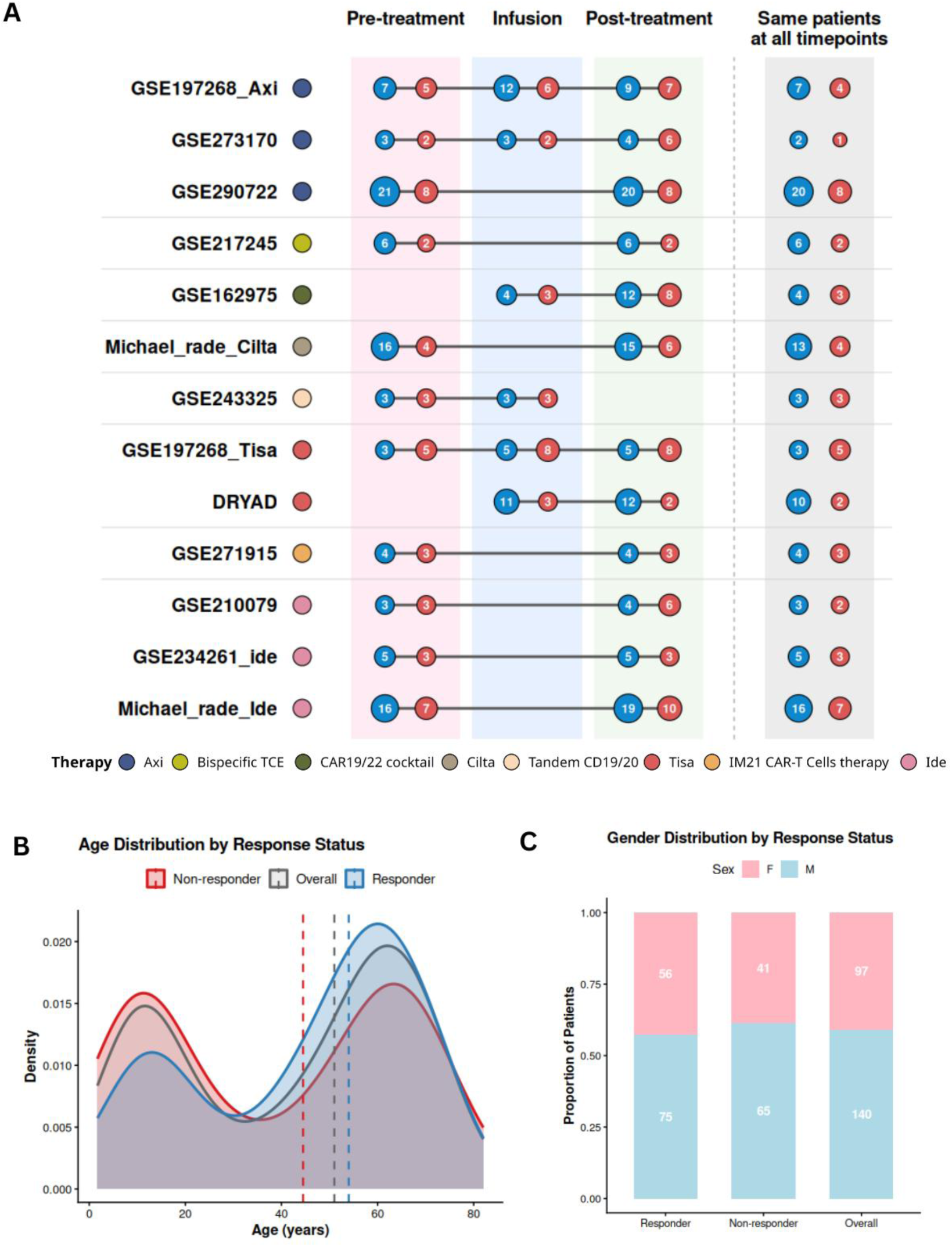
**A)** Longitudinal patient distribution across CAR-T studies. Number of responders (R, blue) and non-responders (NR, red) in studies with longitudinal sampling, stratified by treatment timepoint and CAR-T therapy. The rightmost panel indicates patients represented across all three timepoints. **(B)** Density plot of patient age distribution, stratified by response status (responder (blue) vs. non-responder (red)) and shown for the overall cohort (n = 319). **(C)** Bar plot of patient gender distribution, stratified by response status and shown for the overall cohort (n = 237).

**Figure S2.**
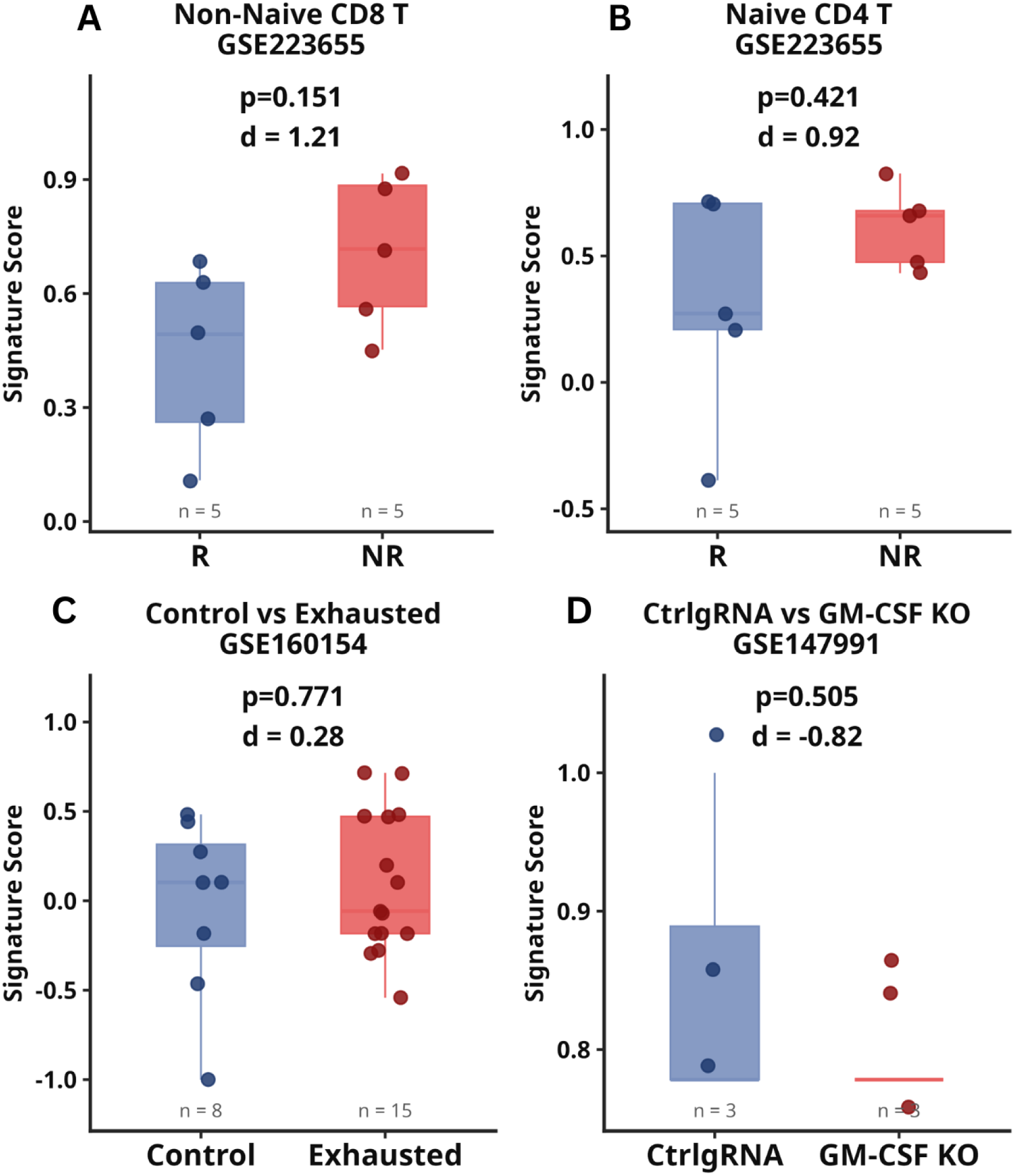
**(A–B)** Pre-treatment apheresis samples (GSE223655) in Non-Naive CD8 T cells (A) and Naive CD4 T cells (B), comparing R and NR. **(C)** In vitro CAR-T exhaustion model (GSE160154); pre-exhaustion control (Day 0, n = 8) versus chronically stimulated exhausted cells (Day 28, n = 15). **(D)** Genetic perturbation model (GSE147991); control gRNA CART19 (NR- like, n = 3) versus GM-CSF KO CART19 (R-like, n = 3). p = Wilcoxon rank-sum test; d = Cohen’s d.

