## Supplementary Methods for "ImmunoCAR Atlas: a pan-immune single-cell resource for dissecting CAR-T therapy response across patient cohorts"

**#Corresponding author**

^#^**Correspondence:**

Dr. Sherry Bhalla

**Supplementary Methods**

**S1. Literature search strategy.**

Searches were performed in PubMed and GEO for studies published up to December 2025, using the following strategies:

1. ("CAR-T" OR "CAR T" OR "chimeric antigen receptor") AND (PBMC OR "peripheral blood mononuclear cells" OR BMMC OR "bone marrow mononuclear cells" OR "infusion product") AND ("single cell" OR scRNA OR scRNA-seq)
2. (acute lymphoblastic leukemia OR diffuse large B-cell lymphoma OR lymphoma OR multiple myeloma) AND (CAR-T OR CD19 CAR OR anti-CD19 CAR-T OR anti-BCMA CAR-T) AND ("single cell" OR scRNA OR scRNA-seq)
3. ("CAR-T resistance" OR non-response OR relapse) AND ("single cell" OR scRNA OR scRNA-seq) AND (PBMC OR BMMC)

Clinical and experimental metadata (response status, tissue source, timepoint, malignancy type, CAR-T target, demographics, MRD status, prior therapy lines, and other reported variables) were manually curated from publications, supplementary materials, and public repositories, and are browsable/downloadable via the "Browse by Study" module of ImmunoCAR Atlas.

**S2. Inclusion criteria.**

Studies were included if they met all of the following: (i) human (*Homo sapiens*) samples; (ii) availability of single-cell transcriptomic data; (iii) reported clinical response information following CAR-T therapy.

**S3. Cell annotation in scRNA-seq data**

For datasets with available raw count matrices, cell type annotation was performed using PopV, an automated cell identity prediction framework that integrates multiple label transfer algorithms and applies consensus voting to assign cell types with associated confidence scores. PopV models were trained using reference atlases from the Tabula Sapiens v2, with tissue-specific references selected according to sample origin. Datasets derived from peripheral blood mononuclear cells (PBMCs) were annotated using the Tabula Sapiens blood reference, whereas datasets derived from bone marrow mononuclear cells (BMMCs) were annotated using the Tabula Sapiens bone marrow reference atlas. Predicted PopV annotations were retained for downstream analyses. For datasets where only normalized expression matrices were available, cell type annotation was performed using CellTypist (v1.6.3). A custom reference model consistent with the PopV annotation framework was applied to ensure annotation comparability across datasets. Final CellTypist predictions and associated confidence scores were retained for downstream analyses.

When available, cell annotations reported in the original studies were preserved alongside automated annotations. (study-reported cell type annotations were available for 18 of the 43 datasets). All downstream analyses were therefore performed using both study-reported labels and standardized automated annotations to enable cross-study comparability while maintaining fidelity to original datasets. PopV annotation, applied uniformly across all 43 datasets, yielded 27 fine-grained cell type categories, these were consolidated into broader lineage-level groups: T cells (CD4+ T cell, CD8+ T cell, naive thymus-derived CD4+ αβ T cell, regulatory T cell, T cell), B cells, plasma cells, precursor cells (hematopoietic cell, hematopoietic precursor cell, hematopoietic stem cell, erythrocyte, common myeloid progenitor, erythroid progenitor cell), neutrophils, basophils, granulocytes, natural killer cells, myeloid cells (monocyte, classical monocyte, macrophage, intermediate monocyte, non-classical monocyte), dendritic cells (myeloid dendritic cell, plasmacytoid dendritic cell), platelets, and fibroblasts.

**S4. T Cell annotation in scRNA-seq data**

To further resolve functional heterogeneity within T cells, we employed two complementary annotation strategies:

1. Marker-based approach using STCAT curated marker gene list
2. Transcriptional program–based approach using StarCAT

For the marker-based annotation, only high-confidence T cells were retained, defined as cells with either a PopV annotation (CD4⁺ T cells, CD8⁺ T) confidence score ≥5 or a CellTypist confidence score ≥0.95. Based on first-level cell type annotations, CD4⁺ and CD8⁺ T cells were analyzed independently. For each subset, batch-corrected latent embeddings were generated using scVI, followed by graph-based clustering to identify transcriptionally distinct subpopulations. Clustering resolutions were set to 0.8 for CD4⁺ T cells and 0.6 for CD8⁺ T cells.

Cluster-specific marker genes were identified by differential expression analysis, retaining the top 1,000 marker genes for each cluster. These marker sets were compared with curated T-cell state signatures obtained from STCAT. Cluster-to-state assignment was performed using two complementary criteria: (i) statistically significant enrichment of cluster marker genes within reference state-specific marker sets (Fisher’s exact test, false discovery rate [FDR] <0.05), and (ii) a minimum overlap of 40% between the cluster marker set and the corresponding reference signature. Clusters satisfying both criteria were assigned to the corresponding functional T-cell state. Clusters that failed to meet these thresholds for any reference state were designated as T unknown cells (heterogenous), representing transcriptionally heterogeneous populations lacking a confident functional assignment.

In parallel, functional annotation was performed using the transcriptional program–based framework StarCAT, which assigns cell states independently of clustering. T-cell populations identified by PopV (CD4⁺ T cells, CD8⁺ T cells, naïve thymic CD4⁺ T cells, and regulatory T cells) were analyzed jointly without prior subdivision. Gene expression matrices extracted from the preprocessed AnnData object were used as input to StarCAT, using the TCAT.V1 reference model. When available, raw count matrices were provided; otherwise, log-normalized expression values were used. StarCAT decomposes single-cell transcriptomes into latent gene expression programs (GEPs) using non-negative matrix factorization and subsequently infers functional T-cell states through reference-based scoring of these programs. Per-cell GEP usage scores and predicted functional state labels were incorporated into the AnnData metadata for downstream analyses.

The marker-based (51 distinct T cell states) and transcriptional program–based annotations (10 distinct T cell states) were evaluated in parallel to provide complementary perspectives on T-cell functional heterogeneity. Whereas the STCAT-based approach identified discrete, cluster-level functional states through canonical marker enrichment, the StarCAT framework inferred continuous, cell-level transcriptional programs independent of clustering, enabling a complementary characterization of T-cell functional diversity.

**S5. Clinical Outcome Harmonization**

Clinical response definitions varied across the included studies and were based on disease-specific evaluation criteria. Reported response categories included complete response (CR), stringent complete response (sCR), partial response (PR), very good partial response (VGPR), stable disease (SD), progressive disease (PD), relapse (RL), long-term response (LtR), minimal residual disease status, prolonged cytopenia status, and progression-free survival (PFS)-based groupings. To enable cross-study comparability, response annotations were harmonized into two standardized categories. The responder group included CR, sCR, PR, VGPR, reported responder status (R), long-term response (LtR), prolonged cytopenia absent, and long PFS group. The non-responder group included reported non-responder status (NR), relapse (RL), PD, MRD-positive SD, prolonged cytopenia present, and short PFS group. Notably, for multiple myeloma (MM) datasets, cases classified as partial response (PR) were categorized as non-responders (NR) rather than responders to ensure consistency with study-specific response definitions and to reflect the underlying disease biology. This classification is also supported by recent literature indicating that PR in MM may represent an insufficient depth of response and is more appropriately grouped with non-responsive disease in comparative analyses. These harmonized response labels were used for all downstream analyses unless otherwise specified.

**S6. scVI-Integrated Atlases**

To enable cross-study comparisons and address the limitations of individual scRNA-seq datasets, we performed data integration across multiple datasets spanning diverse cancer types, therapies, sequencing platforms, and treatment time points. All datasets underwent quality control based on thresholds reported in the original studies, followed by uniform normalization using the scran pooling-based size factor method (v1.22.1) and log1p transformation, which has been shown to perform robustly in integration tasks. For datasets where raw count matrices were unavailable but log1p-normalized expression values were provided, the processed data were retained without additional normalization. Prior to integration, cell-type annotations were harmonized using PopV to ensure consistent labeling across datasets. Integration was performed using the scVI framework (v1.6.6), which leverages deep generative modeling for batch correction and latent representation learning. Raw count matrices were used as input, with patient ID specified as the batch covariate. The model architecture consisted of a negative binomial decoder with a 30-dimensional latent space, 3 hidden layers with 128 nodes, and dropout rate of 0.1. Post-training, batch effects were corrected to eliminate technical biases, latent representations were extracted, and Leiden clustering was applied. These embeddings were visualized via UMAP dimensionality reduction. Integration quality and optimal clustering resolution were evaluated using scIB (v1.1.7). We assessed both biological conservation and batch correction performance using a panel of established metrics. Batch removal was evaluated using graph connectivity, principal component regression (PCR), and batch ASW, while biological conservation was assessed using adjusted Rand index (ARI), normalized mutual information (NMI), cell cycle conservation, isolated label ASW and F1, and cell-type ASW. All evaluation metrics were computed for both the integrated and unintegrated datasets. An overall integration score was calculated as a weighted average of biological conservation metrics (60%) and batch removal metrics (40%). The corresponding atlas-level integration scores and individual evaluation metrics are provided on the Atlas Dashboard page: [Atlas Dashboard – Atlas 1](https://car-t-atlas.igib.res.in/atlas/Atlas%201?utm_source=chatgpt.com).

**S7. T-cell receptor analysis**

Mapping the clonotypic landscape of T cells enables characterization of antigen receptor diversity and identification of clonally expanded populations within the immune compartment. This analysis was performed only for datasets with available single-cell TCR sequencing (scTCR-seq) data. Using single-cell TCR sequencing data, clonotypes were defined based on complementary determining region 3 (CDR3) amino acid sequences of paired TCR α (TRA) and β (TRB) chains. For each cell, a unique identifier was generated by combining sample identity and cellular barcode. Only TRA and TRB chains were retained for analysis. To ensure selection of the dominant receptor sequence, TCR entries were ranked by read count within each cell and chain type, and the highest-abundance CDR3 sequence per chain was selected. Cells lacking a paired TRA and TRB chain were excluded to restrict downstream analyses to fully paired αβ T cells. Clonal frequency was calculated as the proportion of cells sharing an identical paired TRA-TRB clonotype relative to the total number of cells with complete paired TCR information. Clone size was defined as the number of cells belonging to a given clonotype. Based on clone frequency, clonotypes were stratified into expansion groups (hyperexpanded, large, medium, small, and rare) as follows: Hyperexpanded (>10%), large (1–10%), medium (0.1–1%), small (0.01–0.1%), and rare (<0.01%). Clonotype information, including clone identity, clone size, clone frequency, and expansion category, was integrated with the corresponding single-cell transcriptomic dataset via cellular barcodes. This enabled visualization of clonal expansion patterns in low-dimensional embeddings and assessment of clonotype distribution across the annotated T cell population. Furthermore, for each dataset, we compared the distribution of clonotype expansion fractions between responders (R) and non-responders (NR) for each clonotype expansion category. Statistical differences between the two groups were assessed using a two-sided Mann-Whitney U (Wilcoxon rank-sum) test implemented in SciPy (v1.11.4). Comparisons in which either group contained fewer than three patients were considered underpowered and excluded from statistical testing and visualization.

**S8. Cellular composition analysis**

Differences in cell-type composition between clinical response groups were evaluated using scCODA, a Bayesian framework for compositional analysis of single-cell data, implemented via pertpy (v0.6.0). Analyses were performed separately for each tissue source and sampling timepoint (e.g., PBMC_pre, PBMC_post, BMMC_pre, BMMC_post, and infusion product) to account for biological and clinical heterogeneity. Only datasets containing at least two responder and two non-responder samples were included to ensure stable model fitting. Compositional analyses were conducted using multiple annotation hierarchies, including standardized automated annotations (PopV or CellTypist) and study-reported annotations, when available. Clinical response was modeled as a binary variable (responder vs. non-responder). For each tissue/timepoint subset, the number of patients contributing samples was included as a covariate to account for differences in cohort composition. We run scCODA sequentially with default parameters, using No-U-turn sampling (run_nuts_function) and selecting each cell type once as a reference. Within each individual model run, a cell type was considered to have a significant compositional change if its false discovery rate (FDR) calculated as 1 minus the posterior inclusion probability was below the default threshold of 0.05. Credible compositional changes were then determined using a majority-vote framework across model iterations: a cell type was considered credibly associated with response if it met this significance criterion (FDR < 0.05) in more than 30% of model runs. For each cell type, the mean log fold change across iterations was also calculated.

The same scCODA-based approach was applied to the integrated atlas, combining data across studies/cohorts for each tissue/timepoint subset, following an identical modeling strategy as described above (binary response variable, sequential run_nuts_function fitting with each cell type used once as reference, majority-vote credibility threshold, and mean log fold change reporting). The key difference from the single-dataset analyses was in covariate specification: in addition to the number of patients contributing samples, dataset/study of origin was included as a covariate to account for batch effects and technical heterogeneity arising from the integration of multiple independent datasets. This adjustment was intended to reduce the risk of dataset-driven compositional differences being misattributed to clinical response.

Moreover, T cell subsets/states were represented by comparatively low cell numbers within several response-group categories, applying scCODA in this setting risked overfitting or overadjustment given the model's compositional and Bayesian structure. To avoid spurious credible associations under these low-count conditions, T cell composition was instead assessed descriptively using boxplots of per-patient proportions for each T cell state, stratified by response group. This approach was used to visualize overall trends in T cell state distribution between responders and non-responders rather than to test for statistical significance.

**S9. Cell-Cell Interaction Analysis**

To identify intercellular communication patterns associated with CAR-T response, ligand–receptor interaction analysis was performed using CellPhoneDB (v5.0.1), a curated database and statistical framework for inferring biologically relevant cell–cell interactions from single-cell transcriptomic data. This analysis was restricted to individual, sample-level objects and was not performed on integrated or atlas-level objects, as CellPhoneDB requires interactions to be inferred from expression values intrinsic to a single dataset; integration procedures (e.g., batch correction, latent-space embedding) alter the underlying expression matrix in ways that can distort ligand–receptor co-expression signals and are therefore not appropriate for this type of analysis. For datasets with available raw count matrices, expression values were library-size normalized using the *sc.pp.normalize_total()* function in Scanpy (v1.9.3) prior to analysis. Datasets for which only normalized expression matrices were available were used without further normalization. Analyses were conducted separately for each tissue source and sampling timepoint (PBMC_pre, PBMC_post, BMMC_pre, BMMC_post, and infusion product samples) to account for biological and clinical heterogeneity. Interaction inference was performed across multiple annotation hierarchies:

1. standardized automated annotations (PopV or CellTypist), and
2. study-reported annotations when available.

Statistical inference of ligand–receptor interactions was conducted using the CellPhoneDB statistical analysis workflow (*cpdb_statistical_analysis_method.call()*) with default parameters. Ligand–receptor pairs with p-values < 0.05 were considered significant. To compare interaction patterns between responders and non-responders, analyses were restricted to ligand–receptor pairs and interacting cell-type pairs detected in both response groups. For each shared cell-type pair and ligand–receptor interaction, mean interaction scores were compared between response groups. A directional enrichment score was computed by assigning +1 to the group exhibiting a higher mean interaction score for each interaction instance. Scores were aggregated across all shared interactions for a given cell-type pair to generate a cumulative enrichment metric. The final enrichment score was defined as the difference between responder and non-responder cumulative scores, where positive values indicate enrichment in responders and negative values indicate enrichment in non-responders. Statistical significance of differential interaction strength between response groups was further evaluated using the Wilcoxon rank-sum test applied to mean interaction scores for each ligand–receptor pair within a given cell-type pair.

**S10. Patient Cell Composition**

For each dataset, patient-level cell composition profiles were generated by aggregating single cells according to the selected annotation strategy (Consensus PopV (v0.4.2), author-reported, T cell marker-based, or T cell program annotations). Cell counts were quantified for each annotated cell type within individual patient-timepoint groups and normalized by the total number of annotated cells in each group to estimate relative cell-type abundances. These profiles enable longitudinal characterization of immune cell composition across treatment timepoints and tissues at single-patient resolution.

Patient-wise compositional comparison. Differences in cell-type composition between two patients were assessed by constructing a cell type × patient contingency table from the observed cell counts at the selected treatment timepoint. The null hypothesis that cell-type frequencies are independent of patient identity was evaluated using Pearson's χ² test of independence. To identify cell populations contributing to the overall χ² statistic, Pearson residuals were calculated for each cell type as (O−E)/E, where O and E denote the observed and expected cell counts, respectively. Residuals are visualized as a bubble plot, in which bubble size is proportional to the absolute residual magnitude and color denotes the direction of deviation (positive, enrichment; negative, depletion). Cell types with Pearson residuals exceeding |2| were considered the primary contributors to compositional differences between patients.

**S11. Gene Explorer: UMAP visualisation**

For individual datasets, precomputed UMAP coordinates were used as a common low-dimensional representation of the single-cell transcriptomic landscape, with each cell retaining identical coordinates across visualization layers. Gene-specific expression was dynamically overlaid onto this embedding, while a complementary projection was colored by available biological and clinical metadata. The Gene Explorer supports characterization across 15 metadata variables, encompassing cellular identity and state (cell type, T-cell marker- and program-based annotations), patient and dataset identifiers, treatment response, therapy, timepoint, tissue, CAR status, sequencing platform and library preparation, thereby enabling contextual interpretation of gene expression within the cellular and clinical structure of each dataset. For the integrated atlas, precomputed UMAP embeddings provide a unified representation across datasets and are independently annotated using 14 UMAP-level variables, comprising dataset, cancer type, therapy, tissue, timepoint, patient, response, CAR status, sequencing platform, library preparation, study-reported cell type, PopV cell type, and T-cell marker- and program-based annotations. Collectively, these coordinated UMAP projections provide a multidimensional characterization of cellular organization and its association with experimental, clinical, treatment and annotation-related variables across individual datasets and the integrated CAR-T atlas.

**S12. Differential expression analysis**

Differential expression (DE) was performed using two complementary approaches:

1. single-cell limma-trend, to capture cell-level transcriptomic heterogeneity across response groups, and
2. pseudobulk limma-trend, to identify patient-level expression differences while accounting for within-patient cell averaging.

Single-cell gene expression values were obtained from the scVI-normalized layer ("scvi_normalized") and represent library-size normalised counts scaled to 10,000 counts per cell (CPM scale). Prior to statistical modelling, counts were transformed in R by computing log₂(x + 1) to stabilize variance. Genes annotated as mitochondrial (prefix MT-), ribosomal (RPS/RPL), and sex-linked (XIST and Y-chromosome genes) were removed. A per-cell-type expression filter was applied retaining only genes expressed in at least 5% of cells in at least one response group. For each cell type, a design matrix of the form ~0 + Group + patient_id was fitted, where Group encoded the response comparison and patient_id was included as a fixed-effect covariate to account for pseudo-replication from multiple cells per patient. Linear models were fitted with lmFit(), followed by empirical Bayes moderation using eBayes(trend = TRUE, robust = TRUE). Results were extracted with topTable() and reported as logFC, raw p-value (Pvals), and Benjamini–Hochberg FDR (Pvals_adj). A positive logFC denotes greater expression in Group A relative to Group B (e.g., NR vs. R).

To identify consistent patient-level expression differences, pseudobulk profiles were constructed by mean-averaging scVI-normalised expression (layer scvi_normalized_nobatch) across all cells belonging to a given patient within each tissue × timepoint × cell-type stratum, collapsing single-cell resolution into one representative profile per patient. The same gene exclusions were applied (mitochondrial, ribosomal, sex-linked). Strata with fewer than 25 cells or fewer than 2 patients per response group were excluded. The resulting genes × patients matrix was log₂(x + 1)-transformed and genes with zero maximum expression or zero variance were removed. DE was performed using limma-trend (eBayes(trend = TRUE, robust = TRUE)) with design ~ batch + Group when ≥2 batches spanned both response groups, otherwise ~ Group.

**S13. Pathway enrichment analysis**

Over-representation analysis (ORA) of DEGs was performed using the Enrichr API. For single-cell DE results, DEGs were filtered by |logFC| > 0.5 and adjusted p-value < 0.05. For pseudobulk DE results, DEGs were filtered by |logFC| > 0.5 and nominal p-value < 0.05, given the reduced statistical power of patient-level aggregation. Enrichment testing was conducted separately for upregulated and downregulated gene sets against the following GMT libraries: BioCarta_2016, KEGG_2021_Human, MSigDB_Hallmark_2020, and WikiPathways_2024_Human.

**S14. metaCIDER: Cross-cohort differential expression and pathway analysis**

To identify molecular features consistently associated with CAR-T response across independent datasets, metaCIDER integrates dataset-level differential expression and pathway enrichment results within predefined biological strata based on cancer type, CAR-T therapy, tissue, and timepoint. Analyses were performed independently at single-cell and pseudobulk (patient-level) resolutions.

For each dataset, differential expression analysis was performed using limma-trend on scVI-normalised expression values, with the design formula ~ batch + Group for single-cell analysis and the corresponding patient-level design for pseudobulk analysis (as detailed in supplementary methods S12). Dataset-specific DEG tables were generated for each comparison, retaining gene-level log fold-changes (logFC), P-values, and Benjamini–Hochberg-adjusted P-values. These results were subsequently used for cross-cohort gene-level integration and pathway analysis.

For gene-level integration, dataset-specific logFC estimates and corresponding P-values were combined across datasets within each biological stratum using a random-effects meta-analysis. Genes represented in at least two datasets were retained. The meta-analysis procedure, including effect-size estimation, heterogeneity assessment, and multiple-testing correction, is described in S15. In parallel, a frequency-based intersection analysis was performed by applying dataset-level significance thresholds (|logFC| > 0.5 and adjusted *p* < 0.05) and retaining genes meeting these criteria in at least 50% of contributing datasets. Per-dataset logFC and P-values were retained to assess the consistency and direction of gene-level associations.

For pathway-level analysis, enrichment was performed independently for each dataset using hypergeometric over-representation testing against Enrichr-hosted GMT libraries (BioCarta_2016, KEGG_2021_Human, MSigDB_Hallmark_2020, and WikiPathways_2024_Human). Within each dataset, pathways were ranked using a combined score calculated as −log₁₀(FDR) × Gene Ratio, where Gene Ratio represents the number of overlapping genes divided by the total number of genes in the pathway. A cross-dataset frequency filter retained pathways appearing among the top 20 enriched pathways in at least 50% of contributing datasets. For each qualifying pathway, the average −log₁₀(FDR) and average Gene Ratio were calculated across datasets in which the pathway appeared, and their product was used to derive the final Average Rank Score. Pathways enriched in both directions were assigned to the direction with the higher Average Rank Score. The top 10 pathways for each direction were reported, ranked by Average Rank Score and colour-coded according to average −log₁₀(FDR), with the fraction of contributing datasets showing significant enrichment indicated for each pathway.

The same analytical framework was applied independently to single-cell and pseudobulk results, enabling identification of response-associated genes and pathways at both cellular and patient-level resolutions. For pseudobulk-derived gene-level meta-analysis results, nominal pooled P-values rather than BH-adjusted FDR values were used for downstream reporting to preserve sensitivity given the smaller patient numbers.

**S15. metaCIDER: Random-effects meta-analysis**

Gene-level differential expression results from individual datasets were combined using a random-effects meta-analysis to identify transcriptional changes reproducibly associated with CAR-T response across independent cohorts. Analyses were performed separately for single-cell and pseudobulk differential expression results within each predefined biological stratum. Reported log fold-changes (logFC) were used as effect-size estimates, with corresponding standard errors reconstructed from two-sided P-values. Genes represented in at least two datasets were retained for meta-analysis.

Between-dataset variance (τ²) was estimated using restricted maximum likelihood (REML). Pooled logFC estimates and their standard errors were calculated using inverse-variance weighting under the random-effects model, followed by estimation of pooled P-values and 95% confidence intervals. Between-dataset heterogeneity was assessed using Cochran's Q statistic and I², and 95% prediction intervals were calculated to estimate the range of effects expected across comparable independent datasets. Meta-analysis P-values were adjusted across genes using the Benjamini–Hochberg procedure to control the false discovery rate (FDR).

The resulting pooled effect estimates, statistical significance values, and heterogeneity measures were used to characterise the magnitude, direction, and consistency of gene-level associations across cohorts. For pseudobulk-derived meta-analysis results, nominal pooled P-values were used for downstream gene-level reporting rather than BH-adjusted FDR values, given the relatively limited patient numbers.

The analysis was implemented in Python [3.11.15] using NumPy [2.1.3] and pandas [2.2.3] for numerical and tabular operations, SciPy [1.15.1] for REML optimisation and statistical distributions, and statsmodels for multiple-testing correction.

**S16. Pseudobulk matrix construction for Dataset-level signature scoring**

For each dataset, per-patient pseudobulk expression matrices were derived from scVI-normalized, batch-corrected single-cell count matrices stored in the scvi_normalized layer. Prior to aggregation, mitochondrial genes (prefix MT-), ribosomal protein genes (prefixes RPS, RPL), Y-chromosome genes (USP9Y, ZFY, DDX3Y, EIF1AY, KDM5D, NLGN4Y, TSPY1, TXLNGY, UTY, RPS4Y1/2), and XIST-related genes (XIST, TSIX, JPX, FTX) were excluded. Cells were retained only if they carried a valid clinical response annotation (Responder/R or Non-responder/NR) and had tissue and timepoint labels that mapped to the harmonized controlled vocabulary (tissue: PBMC, BM, IP; timepoint: Pre, Post, IP). Pseudobulk samples were defined as unique (patient × tissue × timepoint) combinations; groups with fewer than 25 cells were discarded. Each pseudobulk profile was computed as the arithmetic mean of scVI-normalized expression values across all cells in the group.

**S17. Implementation of the Model Context Protocol (MCP) server for conversational database querying**

Programmatic access to ImmunoCAR Atlas via conversational AI was implemented using the Model Context Protocol (MCP), an open standard that allows large language models such as Claude (Anthropic) to discover and invoke external tools with defined names, descriptions, and input/output schemas. We built a dedicated MCP server (Python, mcp package (<https://py.sdk.modelcontextprotocol.io/>)) as a standalone process, entirely separate from the production Flask application that serves the database; the MCP server never imports, edits, or restarts any file belonging to the existing web application, and interacts with it exclusively through the same read-only REST API already used by the public-facing website, addressed locally to avoid unnecessary external round-trips. The server was configured to listen on an internal port and exposed to Claude over HTTPS via an additive nginx location block layered onto the existing reverse-proxy configuration, leaving the site's primary routing untouched; all configuration changes were syntax-validated and applied via a graceful reload rather than a restart, so live connections to the main site were never interrupted. Because the MCP library enforces host-header validation as a defense against DNS-rebinding attacks, the proxy was configured to present a trusted local host value to the MCP server while forwarding the original public hostname separately, resolving an initial 421 "Invalid Host header" error. Each database capability — dataset listing and metadata, per-dataset differential expression analysis, cell-type proportions, cell-cell interaction analysis, and cross-cohort comparisons — was wrapped as a discrete MCP tool with an explicit schema, so that the language model selects and parameterizes the appropriate tool based on the user's query rather than being given direct database or file-system access. The server runs as a persistent background process with logging, and connectivity was validated end-to-end using both direct local requests and simulated external requests against the public hostname, followed by confirmed successful, independent use by multiple users through their own Claude accounts. No authentication is currently required, consistent with the underlying dataset being publicly available; the integrity of the production application was confirmed unaffected throughout development by comparing file checksums before and after deployment.
